# Inhibition of Nitric Oxide Synthesis by Dexamethasone Increases Survival Rate in *Plasmodium berghei*-Infected Mice

**DOI:** 10.1101/497966

**Authors:** Danilo Reymão Moreira, Ana Carolina Musa Gonçalves Uberti, Antonio Rafael Quadros Gomes, Michelli Erica Souza Ferreira, Rogério Silva Santos, Michael Dean Green, José Ricardo dos Santos Vieira, Maria Fani Dolabela, Sandro Percário

## Abstract

Malaria still presents great epidemiologic importance by its high incidence in the world and potential clinical severity. *Plasmodium* parasites are highly susceptible to changes in the redox balance and the relationship between the redox state of the parasite and host cells is very complex and involves nitric oxide (NO) synthesis. Thus, the present study is aimed at evaluating the effects of NO synthesis on the redox status, parasitemia evolution and survival rate of *Plasmodium berghei*-infected mice. Two-hundred and twenty-five mice were infected with *Plasmodium berghei* and submitted to the stimulation or inhibition of NO synthesis. The stimulation of NO synthesis was performed through the administration of L-arginine, while its inhibition was made by the administration of dexamethasone. Inducible NO synthase (iNOS) inhibition by dexamethasone promoted an increase in the survival rate of *P. berghei*-infected mice and data suggested the participation of oxidative stress in brain as a result of plasmodial infection, as well as the inhibition of brain NO synthesis, which promoted survival rate of almost 90% of the animals until the 15^th^ day of infection, with possible direct interference of ischemia and reperfusion syndrome, as seen by increased levels of uric acid. Inhibition of iNOS caused a decrease of parasitemia and increased survival rate of infected animals, suggesting that the synthesis of NO may stimulate a series of compensatory redox effects that, if overstimulated, may be responsible for the onset of severe forms of malaria.

## INTRODUCTION

Malaria is an acute febrile infectious disease whose etiological agents are protozoa of genus *Plasmodium*. Five species are known to infect man: *Plasmodium vivax, P. falciparum, P. malariae, P. ovale* and *P. knowlesi*. Although there are evidences of the occurrence of the disease since 2700 B.C. (Cox, 2002), it is of epidemiological importance still today by its high incidence in the world and potential clinical severity, causing considerable social and economic losses in the population at risk, especially to ones in precarious conditions of dwelling and sanitation (WHO, 2011).

According to the World Health Organization (WHO) malaria is a significant public health problem in 108 countries and causes approximately 130 million new cases each year (WHO, 2014), resulting in 445 thousand deaths in 2016 (WHO, 2017). About 90% of these deaths occur in sub-Saharan Africa and it is estimated that the disease kills a child every 30 seconds (WHO, 2011).

Usually, the severe cases of the disease are related to the infection by *P. falciparum*. Among the complications that are worth mentioning are cerebral malaria and pulmonary complications (Botelho *et al*. 1996; Van der Heyde *et al*. 2000; Taylor *et al*. 2006; Penet *et al*. 2007). However, the mechanisms that trigger the pathogeny of malaria and the appearance of severe forms are yet not fully elucidated and additional studies are necessary.

In this regard, several authors recently discuss the involvement of free radicals in the physiopathogenesis of malaria (Huber *et al*. 2002; Dondorp *et al*. 2003; Pabon *et al*. 2003; Omodeo-Salé *et al*. 2003; Jaramillo *et al*. 2003; Becker *et al*. 2004; Yazar *et al*. 2004; Wilmanski *et al*. 2005; Kumar and Bandyopadhyay 2005; Dey *et al*. 2009; Percário *et al*. 2012; Vale *et al*. 2015). This involvement can be related to the pathogenic mechanisms triggered by the parasite (Potter *et al*. 2005), as well as by the production of free radicals (Keller *et al*. 2004), and antioxidant defenses (Sohail *et al*. 2007) by host cells as an attempt to fight the infection.

During the development of blood stages of *P. falciparum*, trophozoites increase the viscosity of erythrocytes, by causing modifications on the cell surface that allow its adhesion to the endothelial wall of capillaries, which seems to be a mechanism of defense of the parasite, preventing the passage of parasitized erythrocytes by the spleen and its consequent destruction (Luse and Miller 1971). This cytoadherence phenomenon is mediated by expressed parasite proteins, via stimulation of gene *var*, on the surface of infected red cell, interrupting the blood flow and harming the tissues irrigated by the obstructed vessels (Ferreira *et al*. 2004; Pettersson *et al*. 2005), providing the conditions for the participation of ischemia and reperfusion syndrome (IRS), responsible for free radical production and, consequently, causing oxidative stress (Halliwell and Gutteridge 2015).

In fact, the level of oxidative stress is high in patients infected by *P. vivax*, as detected by the elevation of plasma levels of malondialdehyde - biochemical marker of lipid peroxidation (Farombi *et al*. 2003) - even in patients with non-severe forms of the disease (Pabon *et al*. 2003).

Additionally, oxidative changes in erythrocytes infected with *P. falciparum* seem to be associated to the accelerated aging of these cells and contribute to the development of the anemia displayed by these individuals (Omodeo-Salé *et al*. 2003). The development of anemia can promote changes in the circulatory physiology, leading to the existence of moments of alternate hypoxia and tissue oxygenation at basal levels, thus, another inductor factor of IRS.

Moreover, in response to the infection, activated macrophages and neutrophils act as the natural defense mechanism of the host organism and these generate a large amount of free radicals by activation of respiratory burst, causing an imbalance between the formation of oxidant species and the activity of antioxidants. This imbalance triggers the oxidative stress, being an important mechanism of human host in response to microbial infections that, in the case of malaria, can lead to the death of parasites.

In this regard, the levels of oxidative stress markers in parasitized mice and humans are increased in comparison to non-infected controls (Sohail *et al*. 2007). In these cases, oxidative stress seems to be the result of an increase in free radicals, and not a consequence of the decrease in the levels of antioxidants, reinforcing the suggestion that oxidative stress is an important mechanism induced by the infection (Pabon *et al*. 2003).

In fact, *Plasmodium* is highly susceptible to alterations in the redox balance, which can contribute to clinical manifestations of the severe cases of the disease, such as cerebral malaria (Narsaria *et al*. 2012). In parallel, the relationship between the redox state of the parasite and host cells is very complex and involves production of nitric oxide (NO; Becker *et al*. 2004; Gomes *et al*. 2015).

Nevertheless, the role of NO in malaria is still controversial. Some researchers say that cerebral malaria results from the production of high amounts of NO in order to promote the death of parasites (Favre *et al*. 1999; Maneerat *et al*. 2000), whereas others defend the suggestion that cerebral malaria arises from a low bioavailability of this gas (Gramaglia *et al*. 2006).

Perterson *et al*. (2007), demonstrated that NO synthesis derived from ingested blood in the digestive tract of the mosquito, induces the formation of toxic derivatives, limiting the development of the parasite. As to protect themselves from the damage induced by these toxic nitrogenated derivatives, the mosquito produces pyridoxines, antioxidant enzymes capable of synthetizing NO in response to parasitemia (Herrera-Ortiz *et al*. 2004).

Nahrevania & Dascombe (2006) identified the increase of NO synthesis in *P. berghei*-infected mice and verified its correlation with the increase of the activity of immunologic cells (lymphocytes CD19, macrophages and monocytes).

In fact, some researchers suggest a protective role of nitric oxide in the development of severe malaria and indicate it as possible adjuvant in malaria drug therapy (Yeo *et al*. 2007, 2008; Dhangadamajhi *et al*. 2009). As suggested by Planche *et al*. (2010), the activation of NOS II is essential for the additional production of NO and elimination of the parasite.

On the other hand, Cabrales *et al*. (2011) associated the development of cerebral manifestations of the inadequacy of NO production, which seems to be essential in maintaining cerebral circulatory hemodynamics.

In mice deficient of interleukin 4 (IL-4), it was found that the increase of iNOS expression and the activity of natural killer cells producing IFN-γ, resulted in protection of animals even in the initial phase of infection by *P. berghei* (Saeftel *et al*. 2004).

Moreover, some parasitic molecules are well known as NO inducers, such as the malarial pigment hemozoin, which associated to IFN-γ, is a potent NO inducer in macrophages, involving the kinase regulated extracellular signaling (ERK) pathway and nuclear factor κappa B (NF-κB). It is also known that in the hepatic stage, the defense mechanisms are strictly related to the production of IFN-γ by NK cells, with posterior synthesis of NO (Saeftel *et al*. 2004). In addition, it was found that hemozoin is also responsible for the activation of macrophages by mechanisms partially dependent on NO (Jaramillo *et al*. 2005) and other ROS, such as superoxide (O_2_^−^) and hydrogen peroxide (H_2_O_2_: Brinkmann *et al*. 1984).

Similarly, increased levels of iNOS in human monocytes are associated with non-worsening malaria in patients infected by *P. falciparum* (Chiwakata *et al*. 2000).

Syarifah *et al*. (2003), studying *P. bergh*ei-infected mice susceptible and resistant to the development of cerebral malaria, observed that cytokine expression was increased in resistant animals in relation to susceptible, as well as the expression of NO, worth mentioning the high production of TNF-α in resistant mice, suggesting that the activation of macrophages is significantly greater in those animals.

Three isoforms of nitric oxide synthase enzymes (NOS) were described so far, with two constitutive forms and one inducible form. Constitutive forms produce low amounts of NO for a long period, apparently being responsible for the physiological production of NO. The inducible form (iNOS or NOS II) is activated by factors, such as bacterial lypopolysaccharide (LPS) and cytokines (TNF-α e IFN-γ), producing large amounts of NO in a short space of time (Försterman and Sessa 2012).

All NOS enzymes use L-arginine as substrate, as well as molecular oxygen (O_2)_ and reduced nicotinamide adenine dinucleotide phosphate (NADPH) as co-substrates, and flavine-adenine dinucleotide (FAD), flavine-mononucleotide (FMN) and (6R-)5,6,7,8-tetrahydro-L-biopterine (BH4) as cofactors (Försterman and Sessa 2012). The administration of L-arginine has been employed to stimulate the activity of iNOS in several studies, yet with controversial results (Percário *et al*. 2012).

On the other hand, NOS enzymes can be selectively inhibited. Among the most used inhibitors, N-nitro-L-arginine methyl ester (L-NAME) and N-monomethyl-L-arginine (L-NMMA) inhibit both forms of the enzyme, while aminoguanidine and dexamethasone selectively inhibit iNOS (Walker *et al*. 1997).

Dexamethasone is a glucocorticoid drug and acts on nuclear receptors directly interfering in gene expression in a variety of cell types (Katzung and Trevor 2017) and modulating the transcription of genes involved in the control of inflammatory process (Barnes *et al*. 1993). Since the beginning of the 1990s, some authors identified the effect of dexamethasone inhibiting iNOS expression in most diverse cell types: mesangial cells (Pfeilschifter and Schwarzenbach 1990), murine macrophages (Di Rosa *et al*. 1990), human endothelial cells (Radomski *et al*. 1990), rat hepatocytes (Geller *et al*. 1994), murine fibroblasts (Gilbert and Herschman 1993), and human epithelial cells (Kleinert *et al*. 2004). De Vera *et al*. (1997) attributes this action of dexamethasone by the inhibition of NFκB and to the activation of its inhibitory factor (IFκB). Regardless of the route used by dexamethasone, there is no doubt that NOS inhibition is independent of L-arginine concentration and greatly affects the expression of mRNA for the inducible enzyme (Korhonen *et al*. 2002; Skimming *et al*. 2003).

Administering dexamethasone to *P. berghei* infected mice significantly reduces symptoms of cerebral malaria (Neill and Hunt 1995; Sanni *et al*. 1998)

In the present study it was demonstrated that the selective inhibition of iNOS by dexamethasone reduced the progression of parasitemia in *P. berghei-*infected mice and increased the survival rate of the animals.

## METHODS

Two-hundred and twenty-five male Swiss mice (*Mus musculus*), young adults (25-35 g), from the Evandro Chagas Institute (Belem, PA, Brazil) were randomly divided into three groups, each of them further divided into five sub-groups (n=15 each), according to time of animals’ euthanasia (one, five, ten, fifteen or twenty days after inoculation), and samples of lung tissue and blood were collected for the evaluation of oxidative stress markers, total antioxidant status, uric acid and assessment of percentage of parasitemia, as follows:

**Positive control groups** (N=15 for each sub-group): animals were inoculated with *P. berghei*-infected erythrocytes and received 10 µl of sterile distilled water per 25 g of body weight (gavage) two hours prior to the inoculation of *P. berghei* and daily, until the day of animals’ euthanasia.

**Dexamethasone groups** (N=15 for each sub-group): animals were inoculated with *P. berghei* in the same way that groups PC and treated with dexamethasone, as described below, until the day of animals’ euthanasia.

**L-Arginine groups** (N=15 for each sub-group): animals were inoculated with *P. berghei* in the same way that groups PC and simultaneously treated with L-arginine, as described below, until the day of animals’ euthanasia.

All animals were assigned into sub-groups by simple randomization using the sub-group sequence generated after sortition (Suresh 2011) and were maintained in the vivarium at the Federal University of Pará (UFPA, Belém, PA, Brazil) in polystyrene cages containing five animals each, kept under 12 h light/dark cycles, controlled temperature (25°C), and received rodent chow (Labina™, Presence, Brazil), and tap water *ad libitum* for one, five, ten, fifteen or twenty days after infection and, at the end of each period, animals were submitted to heparin administration (100 UI heparin sulfate, ip.), anesthetized with 50 μl of intraperitoneal ketamine (5%)-xylazine (2%), sample collection, and underwent euthanasia by exsanguination. Absolutely all efforts were made to minimize suffering to animals.

After thoracotomy, blood samples were obtained by cardiac puncture of the right ventricle and both lungs and brain were removed. The project followed the international guidelines for research with experimental animals and procedures were reviewed and approved by the Ethics Committee in Research with Experimental Animals of the Federal University of Pará - CEPAE/UFPA (Report No. MED0126/2013).

### Features of the animal model

Swiss mice are widely used as a malaria model and presents the same pattern of infection progression and basic features of lung and cerebral malarias of other mice species. Moreover, *P. berghei* possesses genomic sequences similar to *P. falciparum* (Otto *et al*. 2014) and cause clinical features on animals that mimic human falciparum malaria (Penet *et al*. 2007). Taken together, the histopathological features described are similar to those displayed in severe malaria human cases.

### Malaria induction

Mice were kept in the vivarium for two weeks and underwent clinical examination prior malaria induction through intraperitoneal inoculation of 10^6^ *P. berghei* ANKA-infected erythrocytes (in 0.2 mL sterile saline solution). The strain of *P. berghei* was supplied by the Neurochemistry Laboratory of the Federal University of Pará - UFPA and three times replicated in Swiss mice before being used in animals of this study.

### Treatments

*Dexamethasone* (Teuto, Cat # 095214): administered in the dose of 5mg/Kg of animal weight.

*L-arginine* (Sigma Aldrich, Cat # A5006): prepared in 0.9% PBS, and administered in a dose of 120 mg/Kg of animal weight (Chatterjee *et al*. 2007).

Both dexamethasone and L-arginine were administered 24h prior infection and every 24h henceforth, until the day of animal euthanasia.

### Tissue processing

After removal, lungs and brain were perfused with PBS to wash out the blood trapped inside. The tissue was weighed and added to PBS in the ratio of 1:10 (m:v). The homogenization process was performed in an ultrasonic cell disruptor (D Cel; Thornton, Indaiatuba, Brazil). During the process, the glass beaker containing the material was kept on ice to prevent sample damage. The homogenate was centrifuged at 175 × *g* (15 min) and the supernatant collected and stored in a freezer at −20°C until analyzed.

### Technical Procedure

Along with blood parasitemia determination, laboratory measurements of trolox equivalent antioxidant capacity (TEAC), thiobarbituric acid reactive substances (TBARS), Uric Acid (AU), and nitrites and nitrates (NN) were performed in duplicate on tissue samples. Internal controls and standards were inserted in each batch for the quality assurance of determinations.

### Determination of parasitemia

*Plasmodium berghei*-infected erythrocytes were counted on blood smears obtained by puncture of the caudal vein of animals on the day of euthanasia (one, five, ten, fifteen, and twenty days of infection). After drying at room temperature, the smear was fixed with methanol for 2 min and stained with Giemsa for 10 min. Subsequently, slides were washed in tap water and, after drying, erythrocytes were counted on an optical microscope (Olympus, CX2) with 100x magnification.

### Determination of Trolox Equivalent Antioxidant Capacity (TEAC)

Trolox (6-hydroxy-2,5,7,8-tetramethylchromane-2-carboxylic acid; Sigma-Aldrich 23881-3) is a powerful antioxidant water-soluble vitamin E analogue. The method proposed by Miller *et al*. (1993) modified by Re *et al*. (1999) was followed, a colorimetric technique based on the reaction between ABTS (2,2’-Azino-bis-3-ethylbenzothiazoline-6-sulfonic acid; Sigma-Aldrich; 1888) with ammonium persulfate potassium (K_2_S_2_O_8_; Sigma-Aldrich; 60490), producing the radical cation ABTS^•^+, chromophore of green/blue color. The addition of antioxidants to ABTS^•^+ reduces it again to ABTS, on a scale dependent on antioxidant capacity, concentration of antioxidants and duration of the reaction. This can be measured by spectrophotometry by observing the change in absorbance read at 734nm for five minutes (Fento, Sao Paulo, Brazil; 800 XI). Finally, the total antioxidant activity of the sample is calculated as its relationship with the reactivity of the Trolox as standard, through the implementation of standard curve under the same conditions.

### Determination of Thiobarbituric Acid Reactive Substances (TBARS)

TBARS is a method that evaluates lipid peroxidation and was used as an indicator of oxidative stress. This technique is based on the reaction of malondialdehyde (MDA), among other substances, with thiobarbituric acid (TBA; Sigma-Aldrich T5500), in low pH and high temperature, yielding MDA-TBA complex of pink color, and absorbance peak at 535 nm.

The technical procedure was performed according to the protocol proposed by Khon and Liversedge (1944), adapted by Percario *et al*. (1994). In brief: initial TBA solution (10 nM) was prepared in phosphate monobasic potassium (KH_2_PO_4_ 75 mM; Synth; 35210) adjusted to pH 2.5 with acetic acid. Two hundred and fifty μL of sample was added to 500 μL of TBA solution, mixed and placed in a water bath (95°C × 60 min); after cooling at room temperature, 2.0 ml of 1-butanol was added, vortex mixed and subsequently centrifuged (175 × *g* × 15 min); 1.0 ml of the supernatant was collected and read at 535 nm (Fento, São Paulo, Brazil; 800 XI). 1,1,3,3, tetraethoxypropane (Sigma-Aldrich; T9889) was used for the implementation of the standard curve.

### Nitrites and nitrates (NN)

Much of nitric oxide released into the bloodstream is swept by hemoglobin in erythrocytes or converted to nitrite (NO_2_^•-^) in the presence of molecular oxygen. Nitrite reacts with oxyhemoglobin, leading to the formation of nitrate (NO_3_^•-^) and methemoglobin. Due to its stability, NO_2_^•-^ has been widely used to confirm the prior existence of NO. The evaluation of this parameter was performed by means of spectrophotometry (Kit Total Nitrite/Nitrate, R & D Systems, KGE001). This technique is based on the quantitative determination of NO, involving the enzyme nitrate reductase, which converts nitrate to nitrite, followed by colorimetric detection of nitrite as a product of pink color, produced by the Griess reaction and that absorbs visible light at 540 nm (PerkinElmer, Victor X3). Nitrite concentration was calculated based on the absorbance found in the nitrites standard curve.

### Uric acid

Performed using the Kit Uric acid UOD-ANA (Labtest, Cat. 51-4/30).

The test is based on the production of H_2_O_2_ from the reaction of uric acid with oxygen and water, catalyzed by uricase. This H_2_O_2_ reacts with acid 3,5-dichloro-3-hydroxybenzene sulphonate (DHBS) and 4-aminoantipyrine in the presence of peroxidase, producing dye antipirylquinonimine. Samples were read at a spectrophotometer at 520nm (Biospectro, SP-22, Brazil).

### Statistical Analysis

Sample size was calculated by the method proposed by Dell *et al*. (2002). The occurrence of discrepant values (*outliers*) was investigated through calculation of interquartile range, which calculates the difference between the third quartile (Q3) and the first quartile (Q1), called dj. Any value lower than Q1 - 3/2 dj or greater than Q3 + 3/2 dj, was considered as *outlier* and, therefore, removed from mathematical calculations.

Aiming at investigating the existence of statistically significant differences between the studied variables between Groups, we applied ANOVA two factors, when the assumption of normality and homoscedasticity was met, or the Mann-Whitney test, when the assumption of normality was not met, which occurred in the case of variable PARASITEMIA. The tests used to access the normality and homoscedasticity of the variables were Kolmogorov-Smirnov and Levene tests, respectively. When the null hypothesis between mean differences between the variables of the study groups was rejected, Tukey’s test was applied, and when a statistically significant difference between medians was detected, Dunn’s test was applied. In addition, within the same group the differences between the initial values (1 day of infection) and late values (20 days of infection) were studied by the Student’s unpaired t test.

The existence of correlation between the variables was also analyzed by Pearson’s correlation coefficient, considering all points obtained separately for each group studied. For the statistically significant correlations, intensities were assigned as follows: r up to 0.30 (r < 0.30) as weak correlation; r between 0.31 and 0.70 (0.31 < r < 0.70), as moderate correlation; r between 0.71 and 1.00 (0.71 < r < 1.00) as strong correlation.

For the purposes of tests ANOVA and Mann-Whitney, statistical package SigmaStat version 3.5 was used, whereas for the calculation of correlations the statistical package SPSS version 17.0 was used. All statistical tests were applied considering the significance level of 5% (*p*<0.05).

### Availability of Data and Materials

Data and full description of methods and materials are available at Zenodo repository, at https://zenodo.org/record/45202#.VqeqifLVyid.

## RESULTS

As expected, parasitemia of infected animals progressively evolved in all groups, but the rate of progression was lower in dexamethasone-treated animals, which presented lower values than the other two groups at the end of the period of 20 days (p=2.8×10^−5^ *vs*. L-arginine and p=0.0227 *vs*. control; Fig. 1). L-arginine-treated animals presented numerically higher values than the control group, but without statistical significance (p=0.3048).

**Figure 1.**
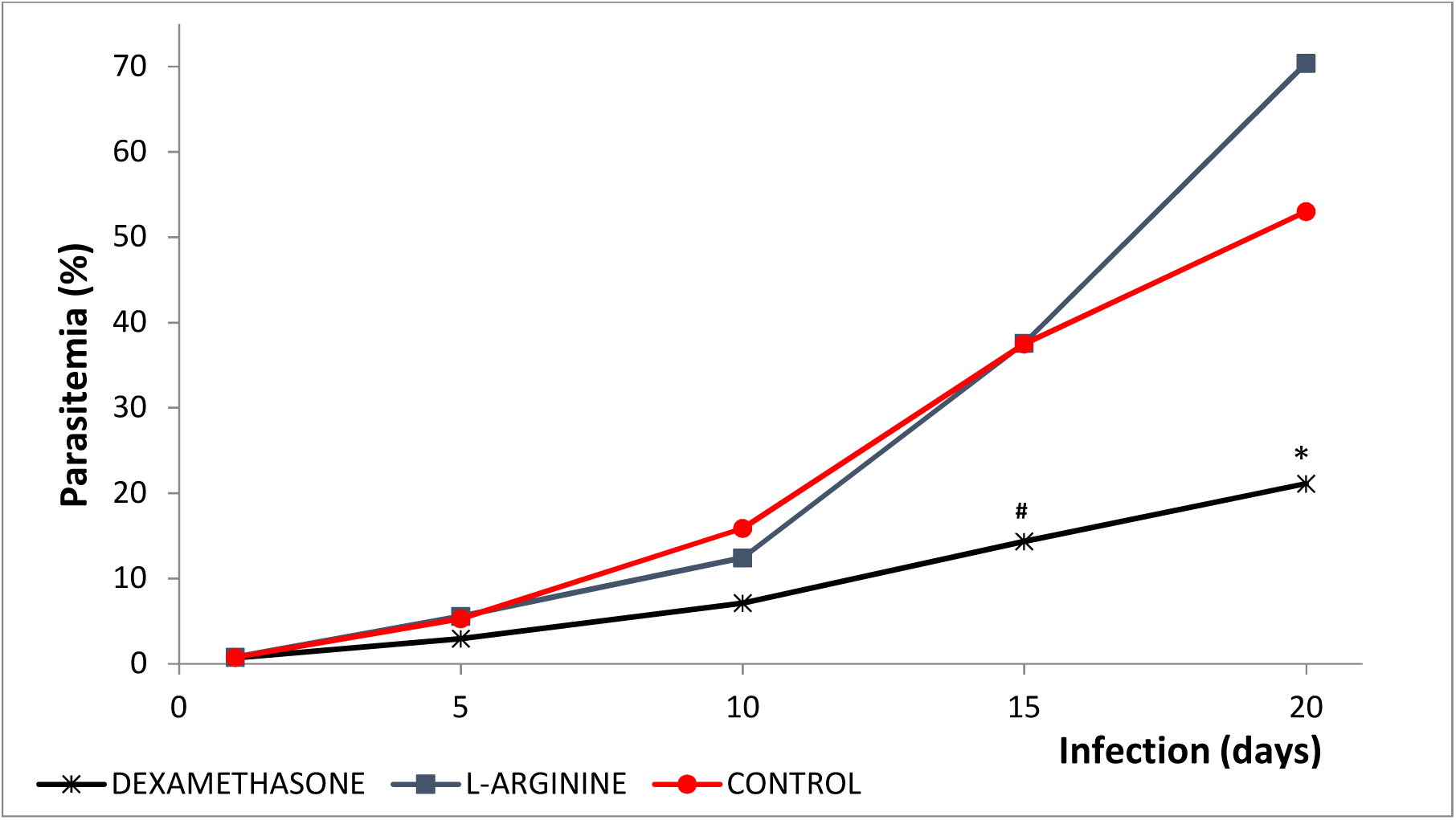
Progression of parasitemia in *Plasmodium berghei*-infected Swiss mice. Animals were pre-treated and received a daily dose of DEXAMETHASONE, L-ARGININE, or PBS (CONTROL). ^#^ p=6.8×10^−6^ *versus* L-ARGININE and p= 7.3×10^−5^ *versus* CONTROL; * p=2.8×10^−5^ *versus* L-ARGININE and p=0.0227 *versus* CONTROL.

Similarly, the survival rate of dexamethasone-treated animals was significantly greater than that of the other groups, which behaved in a similar way, with 60% of animals alive at the end of the period of 20 days of infection (Fig. 2).

**Figure 2.**
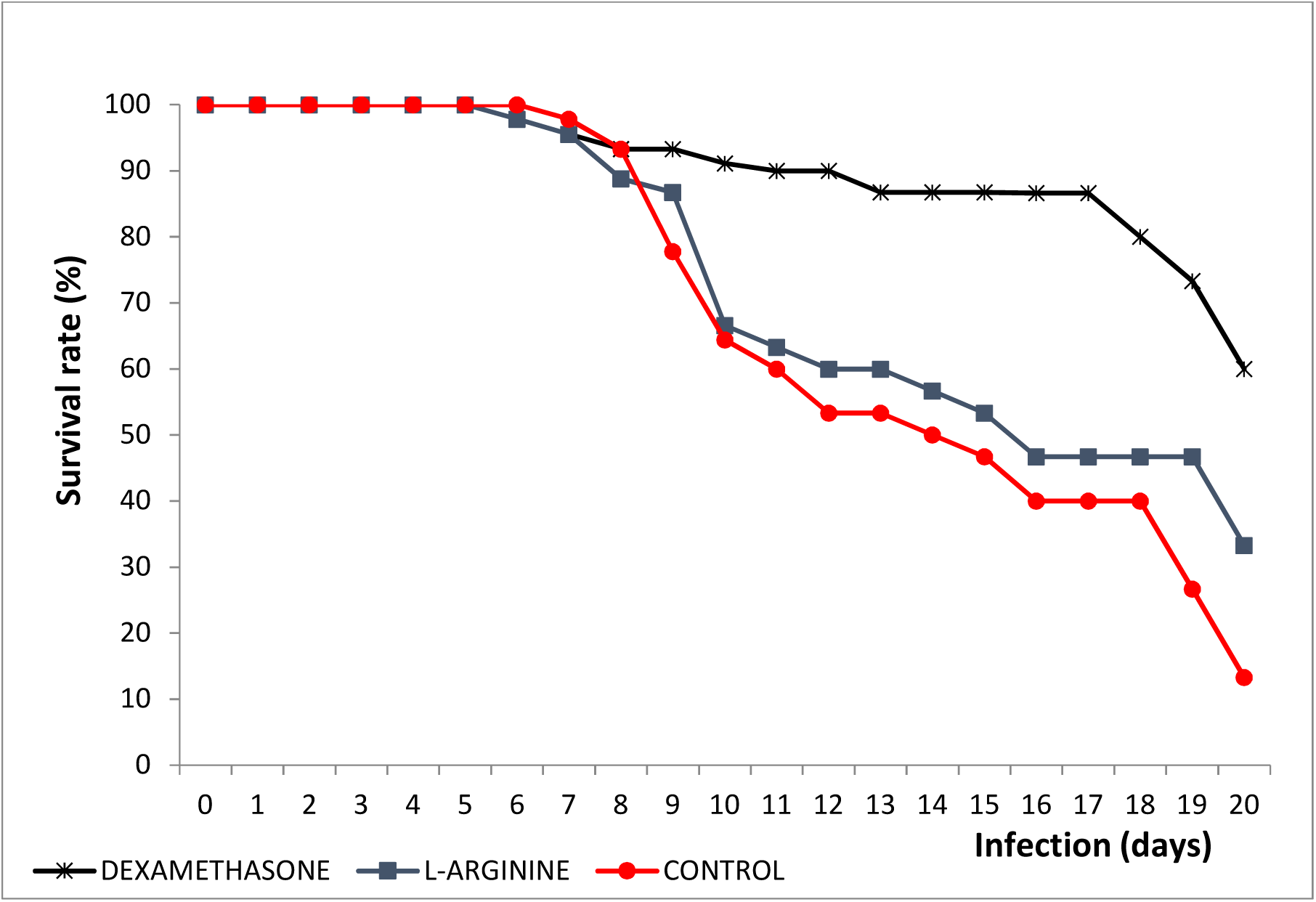
Survival rate of *Plasmodium berghei*-infected Swiss mice. Animals were pre-treated and received a daily dose of DEXAMETHASONE, L-ARGININE, or PBS (CONTROL).

## TEAC

For the lung samples, all groups showed a slight decrease of TEAC values along the period of infection, however without statistically significant differences (Fig. 3A). Nevertheless, at the end of 20 days of infection, the group of animals treated with dexamethasone presented statistically lower values than the other two groups (p=0.0281 *vs*. L-ARGININE and p=0.0033 *vs*. CONTROL). For brain samples a similar behavior was observed, however an important decrease of TEAC after 10 days of infection was identified (1 day *vs*. 10 days, p=0.0009 for L-ARGININE and p=7×10^−6^ for DEXAMETHASONE), with both treated groups presenting values lower than the control group (p=0.0360 *vs*. L-ARGININE and p=0.0261 *vs*. DEXAMETHASONE). However, after the 10^th^ day of infection the DEXAMETHASONE group presented an increase in TEAC values, displaying statistically higher values than the other groups (p=0.0357 *vs*. L-ARGININE and p= 0.0005 *vs*. CONTROL).

**Figure 3.**
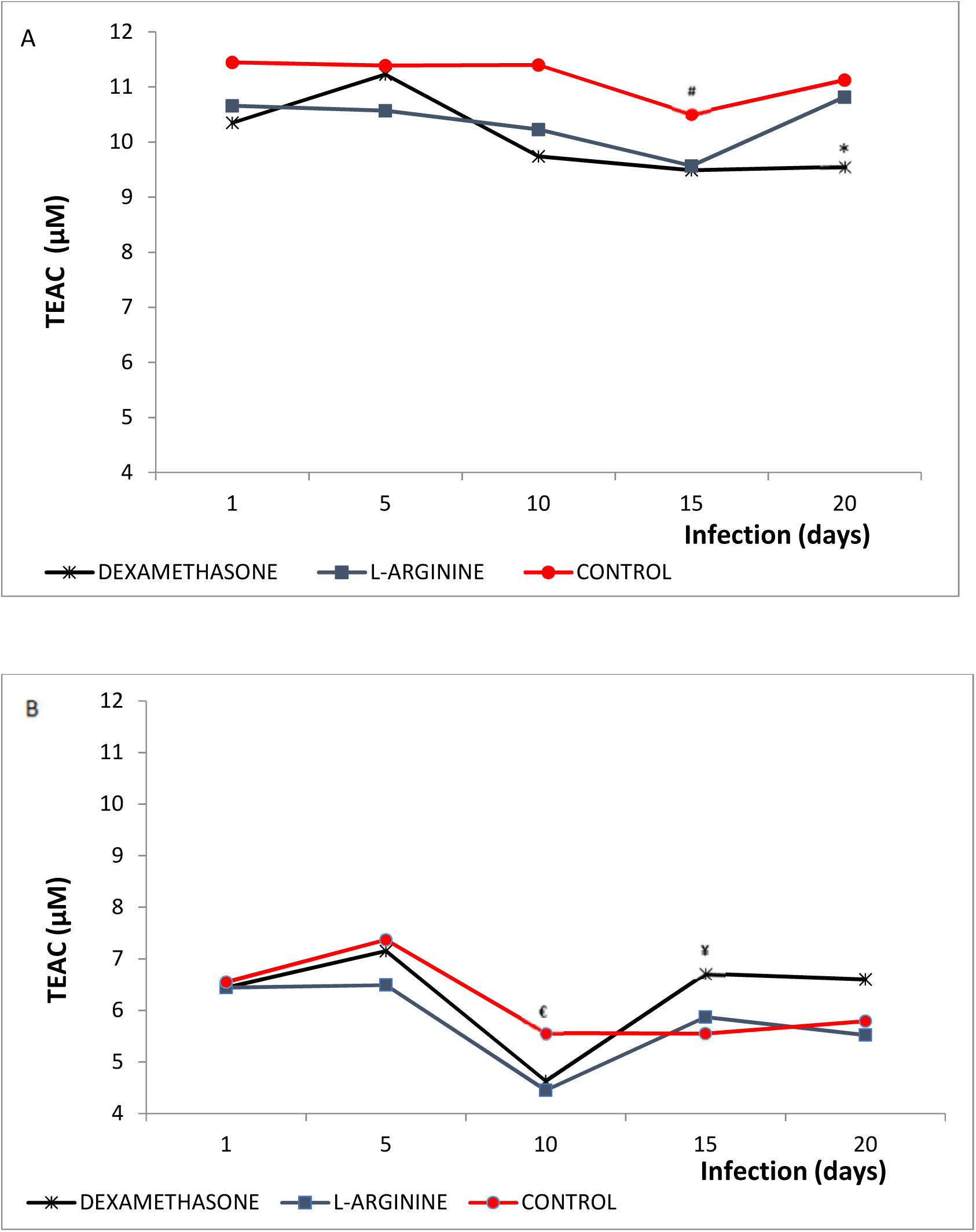
Trolox Equivalent Antioxidant Capacity (TEAC) in lungs (A) and brains (B) of *Plasmodium berghei*-infected Swiss mice. Animals were pre-treated and received a daily dose of DEXAMETHASONE, L-ARGININE, or PBS (CONTROL). **^#^** p=0.0401 *versus* DEXAMETHASONE; **\*** p=0.0281 *versus* L-ARGININE and p=0.0033 *versus* CONTROL; **^€^** p=0.0360 *versus* L-ARGININE and p=0.0261 *versus* DEXAMETHASONE; **^*¥*^** p=0.0357 *versus* L-ARGININE and p=0.0005 *versus* CONTROL.

## TBARS

Although none of the groups have presented important variation during the period of infection, Group L-ARGININE presented higher pulmonary TBARS values than the DEXAMETHASONE group at the end of the experiment (p=0.0282). On the other hand, for brain samples, Group L-ARGININE showed progressive evolution over the period of the infection, with higher TBARS values in the 20^th^ day of infection in relation to the first day (p=4.7× 10^−7^), but with no differences in relation to the other groups (Fig. 4).

**Figure 4.**
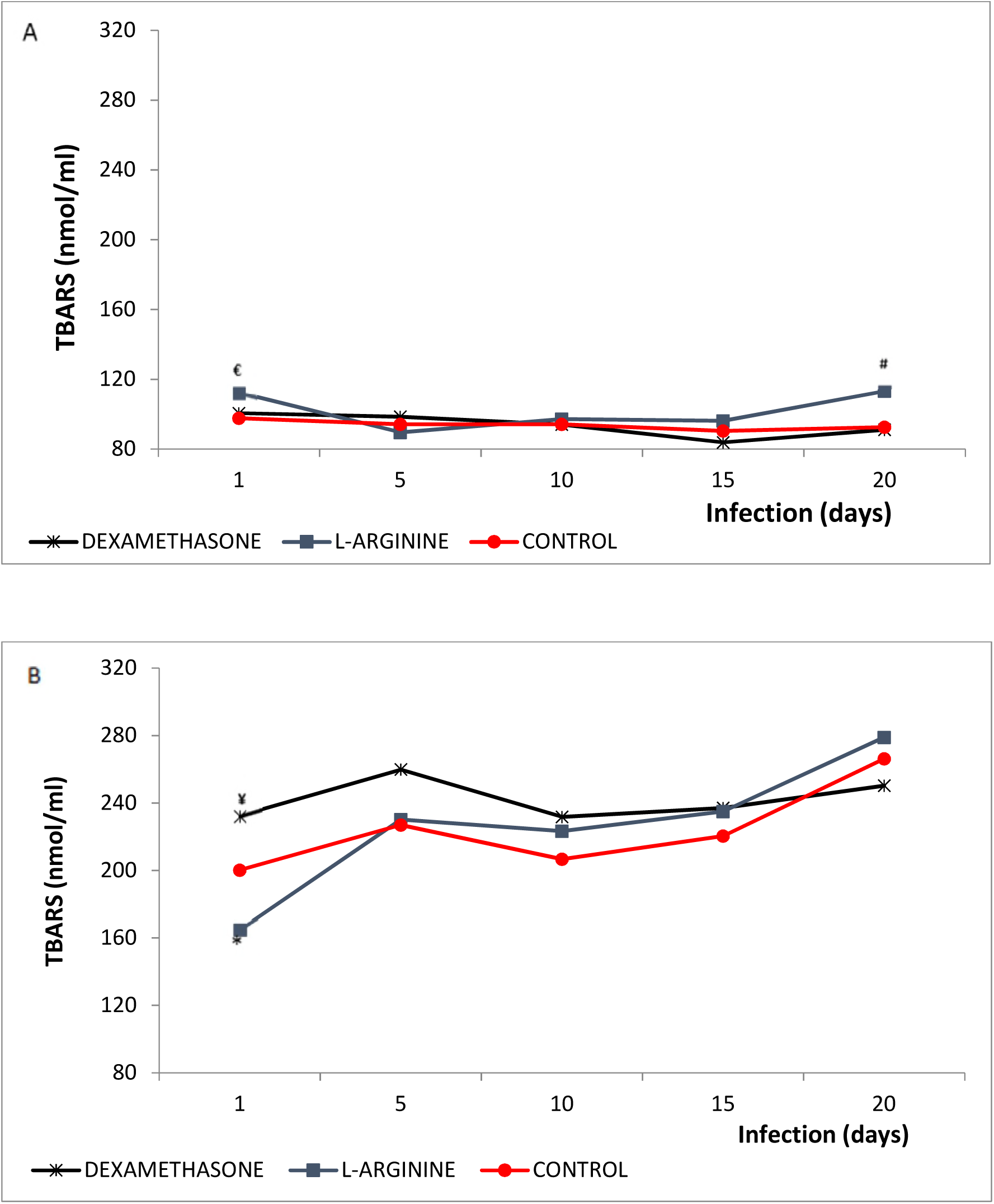
Thiobarbituric Acid Reactive Substances (TBARS) in lungs (A) and brains (B) of *Plasmodium berghei*-infected Swiss mice. Animals were pre-treated and received a daily dose of DEXAMETHASONE, L-ARGININE, or PBS (CONTROL). **^€^** p=0.0294 *versus* CONTROL; **^#^** p=0.0282 *versus* DEXAMETHASONE; **\*** p=0.0005 *versus* DEXAMETHASONE and p=0.0029 *versus* CONTROL; **^*¥*^** p=0.0060 *versus* CONTROL.

The collective analysis of the values of TEAC and TBARS shows a quite unique pattern: while for lung samples the values of TEAC obtained are found in a high range of absolute values (9-12 µM), brain samples are in a low range (4-8µM), whereas TBARS values presenting opposing behavior, i.e. for lung samples the values are in low range (80-120nmol/mL) and brain samples in high range (160-280nmol/mL).

### Nitrites and nitrates

No significant differences in the evolution of NN levels in any of the groups throughout the period of infection were seen, nor between groups for the lung samples (Fig. 5). However, for brain samples, group DEXAMETHASONE presented lower values than the other two groups during the studied period, culminating with statistically significant differences in the 20^th^ day (p=0.0058 *vs*. L-ARGININE and p=0.0201 *vs*. CONTROL).

**Figure 5.**
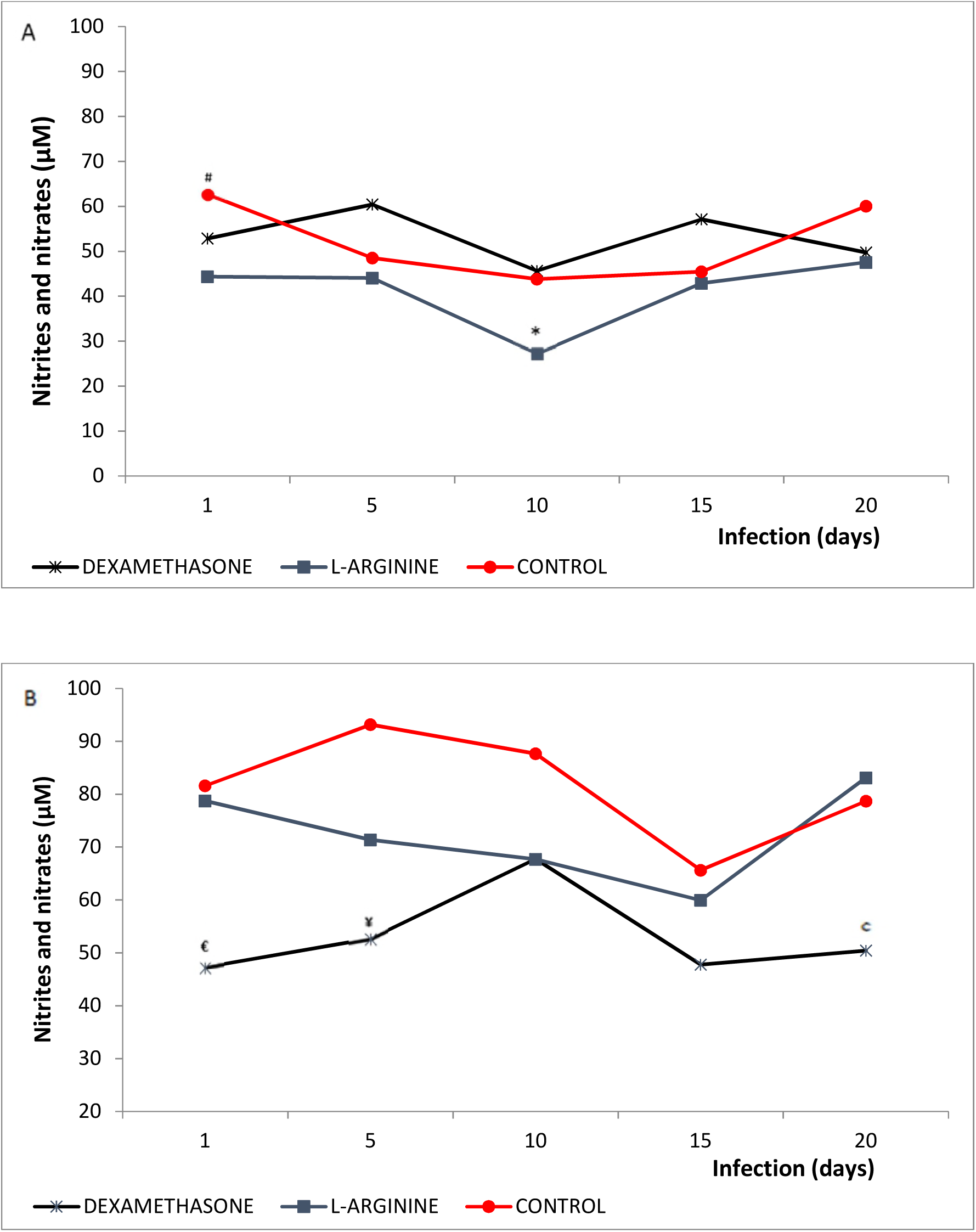
Nitrites and Nitrates in lungs (A) and brains (B) of *Plasmodium berghei*-infected Swiss mice. Animals were pre-treated and received a daily dose of DEXAMETHASONE, L-ARGININE, or PBS (CONTROL). **^#^** p=0.0005 *versus* L-ARGININE and p=0.0394 *versus* DEXAMETHASONE; **\*** p=3.1×10^−5^ *versus* DEXAMETHASONE and p=1.5×10^−4^ *versus* CONTROL; **^€^** p=3.4×10^−6^ *versus* L-ARGININE and p=5.0×10^−4^ *versus* CONTROL; **^*¥*^**p=3.6×10^−4^ *versus* L-ARGININE and p=4.6×10^−4^ *versus* CONTROL; **^c^** p=0.0058 *versus* L-ARGININE and p=0.0201 *versus* CONTROL.

### Uric acid (AU)

No temporal variation in AU values for lung samples in any of the groups were found (Fig. 6). However, Group L-ARGININE presented lower values than the other two groups from the first to the 15^th^ day of infection. Similarly, for brain samples, group L-ARGININE presented lower values than the other groups, with statistical significance at the 20^th^ day of infection (p=0.0395 *vs*. CONTROL and p=0.0407 *vs*. DEXAMETHASONE). In contrast, group DEXAMETHASONE presented progressive behavior over the infection time, with brain AU values significantly greater for the 20^th^ day in comparison to the first day (p=3.9×10^−4^). Another noteworthy observation is that pulmonary AU values stood in a higher range (40- 140mg/dL) than the brain (10-55mg/dL) for all groups.

**Figure 6.**
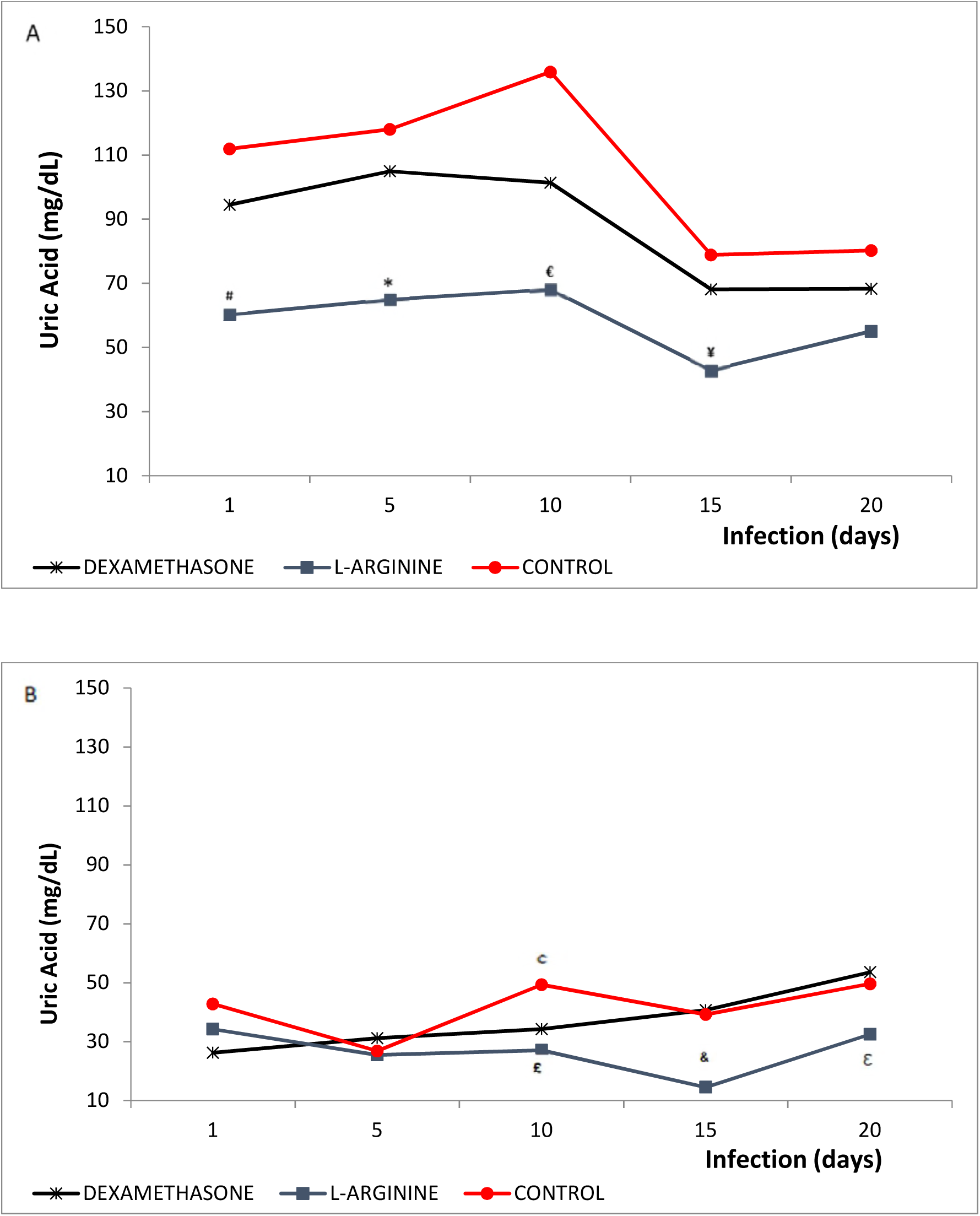
Uric Acid levels in lungs (A) and brains (B) of *Plasmodium berghei*-infected Swiss mice. Animals were pre-treated and received a daily dose of DEXAMETHASONE, L-ARGININE, or PBS (CONTROL). **^#^** p=0.0033 *versus* DEXAMETHASONE and p=4.3×10^−6^*versus* CONTROL; **\*** p=0.00058 *versus* CONTROL; **^€^** p=0.00095 *versus* CONTROL and p=0.00054 *versus* DEXAMETHASONE; **^*¥*^** p=0.00071 *versus* CONTROL and p=0.01167 *versus* DEXAMETHASONE; **^c^** p=0.0080 *versus* DEXAMETHASONE and p=0.0024 *versus* L-ARGININE; **^£^** p=0.0479 *versus* DEXAMETHASONE; ^**&**^ p=0.0029 *versus* CONTROL and p=0.0001 *versus* DEXAMETHASONE; **^ε^** p=0.0395 *versus* CONTROL and p=0.0407 *versus* DEXAMETHASONE.

### Correlation studies

#### PARASITEMIA vs. TBARS

The correlation between TBARS and PARASITEMIA revealed the existence of a negative and significant correlation only for the lung samples from the group DEXAMETHASONE (Additional file 1 Fig. 1; r=-0.29; p=0.026). The CONTROL group presented a negative correlation, however without statistical significance (r=- 0.10; p=0.20), while for group L-ARGININE this correlation showed positive but non-significant values (r=0.06; p=0.673). For brain samples a positive trend was observed for all groups, but only with significance for group L-ARGININE (Additional file 1 Fig. 2; r=0.46; p=0.002).

#### TBARS vs. URIC ACID

A positive correlation was observed for these parameters in both samples and for groups CONTROL (Additional file 1 Fig. 3-4; r=0.33 and p=0.02, for lung; r=0.45 and p=0.050, for brain), and DEXAMETHASONE (r=0.26 and p=0.041, for lung; r=0.28 and p=0.045, for brain). For group L-ARGININE, in both samples, the values of the coefficient of correlation approached zero (r=0.08 and p=0.140, for lung; r=0.03 and p=0.858, for brain).

#### NN vs. TBARS

For lung samples the existence of significant correlation for any of the studied groups was not observed (Additional file 1 Fig. 5). However, for brain samples, both groups CONTROL and DEXAMETHASONE presented significant positive correlations (Additional file 1 Fig. 6; r=0.30 and p=0.048 and r=0.34 and p=0.014, respectively).

#### TEAC vs. TBARS

The existence of positive correlation in both samples and for both groups CONTROL (Additional file1 Fig. 7-8; r=0.32 and p=0.024, for lung; r=0.50 and p=0.009, for brain) and DEXAMETHASONE, (r=0.23 and p=0.031, for lung; r=0.27 and p=0.050, for brain) was seen. For the group L-ARGININE, in both samples, the values of the coefficient of correlation were negligible (r=0.05 and p=0.821, for lung; r=0.14 and p=0.374, for brain).

#### NN vs. PARASITEMIA

No significant correlation was found for any of the groups, nor for any of the samples studied (Additional file1 Fig. 9-10).

#### Other Correlations

In addition to the studies of correlation presented, we tested the following correlations: TEAC *vs*. NN (Additional file 1 Fig. 11-12); TEAC *vs*. URIC ACID (Additional file 1 Fig. 13-14), TEAC *vs*. PARASITEMIA (Additional file 1 Fig. 15-16), NN *vs*. URIC ACID (Additional file 1 Fig. 17-18), and URIC ACID *vs*. PARASITEMIA (Additional file 1 Fig. 19-20).

## DISCUSSION

Malaria is a disease of high incidence worldwide and infection by *P. falciparum* are responsible for severe manifestations of the disease and the majority of the cases of deaths related to this disease. Cerebral malaria and pulmonary complications are among the noteworthy complications of malaria and are similar to the clinical manifestations associated with *P. berghei* infection on the animal model employed in the present study (Botelho *et al*. 1996; van der Heyde *et al*. 2000; Taylor *et al*. 2006; Penet *et al*. 2007). The mechanisms that trigger the pathogeny of malaria and the appearance of severe forms are not fully elucidated yet. In this context, oxidative stress, through NO synthesis, seems to play a dubious but important role. Nevertheless, survival rate results pointed to a significant increase in the percentage of survival for the groups of mice treated with the iNOS inhibitor dexamethasone in comparison to other groups (Fig. 2), which seems to be correlated with the evolution of parasitemia in these animals, which has changed very little in this group and remained, from the 5^th^ day henceforth significantly lower than the other groups (Fig. 1).

It is important to highlight that dexamethasone is a non-steroid anti-inflammatory drug that acts through iNOS mRNA synthesis inhibition (Korhonen *et al*. 2002). In this sense, it is possible that NO acts oxidatively, both inducing the worsening of the disease, as favoring the increasing of parasitemia.

Therefore, the effect of dexamethasone on the evolution of the parasitemia can promote inhibition of oxidative stress, as may be suggested by the existence of a negative correlation between TBARS and PARASITEMIA found only for the animals of group dexamethasone (Additional file Fig. 1). In the same way, the effect of L-arginine is consistent with the effect found in this correlation for the animals of Group L-ARGININE, where there is the reversal of this pattern, presenting positive values of correlation.

Contrary to that mentioned by some authors (Yeo *et al*. 2007, 2008; Dhangadamajhi *et al*. 2009; Planche *et al*. 2010; Cabrales *et al*. 2011) that attribute a protective role to nitric oxide in malaria, mice treated with L-arginine remained with percentage of parasitemia and survival rate comparable to group CONTROL (Fig.1-2), suggesting that NO synthesis is not involved among the initial mechanisms of host defense and, therefore, may not contribute to the elimination of parasites. However, the mentioned studies have measured the survival rate of animals treated with Dipropylene triamine NONOate, a natural donor of nitric oxide, active in acid PH (common in malaria) whose action, unlike L-arginine, is independent of enzyme activation.

### Pulmonary findings

The mechanisms responsible for triggering the syndrome of respiratory anxiety displayed in malaria patients are multifactorial. However, according to some authors there are no doubts about the participation of free radicals (Gachot *et al*. 1995; Taylor *et al*. 2006; Gillrie *et al*. 2007), which directly affect the cell membranes, attacking the endothelium and changing vascular permeability.

Among the free radicals involved in this process, NO seems to play an important role. However, the paradox of the actuation of this molecule in pulmonary complications is evident: while some authors suggest that the inhalation of this gas is a potential treatment of these complications (Rabkin *et al*. 2001; Schreiber *et al*. 2003; McClintock *et al*. 2007; ter Horst *et al*. 2007), others blame nitric oxide synthesis as responsible for causing the respiratory distress syndrome (Adhikari *et al*. 2007), in particular as a result of the activation of iNOS (Mikawa *et al*. 2003; Baron *et al*. 2004).

Additionally, superoxide radical, stimulated by substances derived from the inflammatory process, such as TNF-α, IL-1α and lipopolysaccharide (through the increase on NADPH-oxidase activity), are also present in the syndrome, which also seems to be related with the increase of nitric oxide synthesis (Muzaffar *et al*. 2003). However, the reaction between these two free radicals (nitric oxide and superoxide) is known to produce a potent third free radical, peroxynitrite (Katzung and Trevor 2017). Thus, nitric oxide seems to play a fundamental role, once that iNOS activation is associated with the induction of NADPH-oxidase and the production of peroxynitrite (Muzaffar *et al*. 2003; Adhikari *et al*. 2007).

As a defense mechanism against the cellular damage caused by oxidative stress, there is an increase in the production of antioxidant molecules both from alveolar surfactant production (which is notoriously hyper secreted), as well as by the increase in antioxidant enzyme activity (Rahman and MacNee 2000; Oury *et al*. 2002; Christofidou-Solomidou *et al*. 2003; Kinsella *et al*. 2005; Saxena *et al*. 2005; Bein *et al*. 2009).

In the present study there was no significant variation in the dosages of TEAC and TBARS levels in the control group along the infection period (Fig. 3-4). However, it was observed the occurrence of a positive correlation between these parameters for both samples tested (Additional file Fig. 7-8), suggesting that the increase in oxidative stress resulting from the infection induced the increase of antioxidant defenses, but could not be reversed by it.

Additionally, the behavior of the TBARS *vs*. TEAC correlations and TBARS *vs*. Uric acid was similar (Additional file Fig. 3-4), suggesting that uric acid is an important component of the antioxidant defense of these animals, or IRS is associated with the infection. In this sense, the absence of correlation between these parameters found in Group L-ARGININE for both samples is further evidence of the absence of the IRS in animals in this group, probably as a result of vasodynamic effects attributable to NO.

Among the treatments, the only one that showed significant correlation between TBARS and PARASITEMIA, was dexamethasone (r=-0.29, p=0.026; Additional file Fig. 1) and can suggest that the selective inhibition of iNOS, associated to the anti-inflammatory potential of dexamethasone, decrease the lipid peroxidation even with the increase of parasitemia. This suggestion is reinforced by the finding of negative correlations between TEAC *vs*. PARASITEMIA and URIC ACID *vs*. PARASITEMIA (Additional file Fig.15 and 19), since enzymatic antioxidant defenses and IRS suffer direct influence of lipid peroxidation.

In this experimental model, considering that all animals were exposed to the same food supply, high values of uric acid indicate the existence of ischemia and reperfusion syndrome (Halliwell and Gutteridge 2015), and may be caused by the decrease of the caliber of blood vessels, by anemia, or by obstruction of the blood flow by the occurrence of cytoadherence.

It was found that a significant positive correlation for URIC ACID and TBARS levels in both samples and for both groups CONTROL and DEXAMETHASONE, suggesting that IRS arises from the increased oxidative stress in these animals as a consequence of disease progression, as well as that NO synthesis may not exert important effect in this case. On the other hand, for group L-ARGININE, in both samples, the values of the coefficient of correlation approached zero, suggesting adequate blood supply to these tissues, possibly as a result of NO-attributable vasodilation.

The treatment with L-arginine did not promote any modification in the antioxidant capacity during the period studied (Fig. 3). On the other hand, it has significantly increased lipid peroxidation, but only in the first day of infection (Fig. 4A). The decrease in lipid peroxidation in subsequent days can be explained by the decrease in the IRS, justified by the low levels of uric acid for animals of this group during the entire period of infection (Fig. 6A).

The high correlations (moderate to strong) for TEAC *vs*. URIC ACID in all groups (Additional file Fig. 13) arises from the simple fact that uric acid, by itself, is an antioxidant, in addition of being a marker of IRS. The same is true for the positive correlations between TEAC *vs*. NN (Additional file Fig.11) and NN *vs*. URIC ACID (Additional file Fig. 17), displayed by most of the groups.

Among the most unusual results, it is noteworthy the absence of differences in the levels of pulmonary nitrites and nitrates, independent of the use of inhibitor (dexamethasone) or stimulator of their synthesis (L-arginine; Fig. 5A). The possible explanations for such phenomena arising out of compensatory physiological effects, such as vasoconstriction caused by the NOS inhibition, which seems to stimulate the production of mediators that cause vasodilation such as acetylcholine and bradykinin, which are bronchoconstrictors nonetheless (Silverthorn 2010). Conversely, it is possible that pulmonary hypertension on malaria, reported by Lacerda *et al*. (2009), as caused by the inhibition of NO by treatment with dexamethasone, along with the need of oxygen as a result of hemolysis, stimulates the synthesis of eNOS, which increases the expression of eNOS receptors in the lungs (Beleslin-Čoki *et al*. 2011).

The opposite effect happened for group L-ARGININE, in which it was expected an increase in nitrites and nitrates, but despite the lack of statistical significance, stood numerically below of the other two groups. It is worth mentioning that after formed, L-arginine can follow two paths: the formation of ornithine and urea (action of arginase) or the formation of citrulline and NO (action of NOS). Additionally, interleukins (IL) 13 and 14 act over arginase directing L-arginine to the synthesis of ornithine that is converted, by the action of an aminotransferase, to proline. This route has fibrogenic role, since proline is an essential amino acid in collagen (Lee *et al*. 2001). On the other hand, cytokines interferon γ (IFN-γ), tumor necrosis factor-α (TNF-α) and IL-12, optimize the formation of NO and citrulline from the action of iNOS over L-arginine (Hesse *et al*. 2000). Thus, it is likely that the excess of L-arginine, depending on the profile of cellular response stimulated, follow the arginase route, promoting the clearance of pulmonary nitrites and nitrates (Modolell *et al*. 1995; Chiaramonte *et al*. 1999; Hesse *et al*. 2000; Lee *et al*. 2001), resulting in fibrinogen synthesis, in an attempt to revert pulmonary damage caused by oxidative stress.

Another possibility is that the vasodilation produced by NO excess increase availability of O_2_, substrate of NADPH oxidase, resulting in greater production of superoxide radical and, consequently, of peroxynitrite (Muzaffar *et al*. 2003; Adhikari *et al*. 2007). According to Wedgwood *et al*. (2012), peroxynitrite levels impose a negative feed-back on NOS, i.e., the more peroxynitrite is synthesized, greater inhibition of NOS.

Additionally, the absence of differences between the groups for the values of pulmonary NN may be the result of the existence of a complex system of non-adrenergic non-cholinergic (NANC) neural fibers in the lungs of mammals, capable of producing large quantities of NO (Gaston *et al*. 1994) and, therefore, to masque NO levels arising from malaria in this tissue. This suggestion is reinforced by the absence of correlation between NN and TBARS levels in all groups for lung samples (Additional file Fig. 5). In contrast, for brain samples, both groups CONTROL and DEXAMETHASONE showed significant positive correlations, while group L-ARGININE showed no correlation between these parameters (Additional file Fig. 6). These data suggest that, at least partially, oxidative stress associated with the development of the disease is derived from the production of NO, as pointed out by several authors, in addition to the participation of IRS (Percário *et al*. 2012), which may have been reversed in the animals treated with L-arginine, due to its vasodilator effect.

### Cerebral findings

Similar to the pulmonary features of the disease, cerebral edema seems to determine the pathological onset of severe malaria. However, the increase of intracranial pressure due to cerebral edema results in greater risk of death. This abnormality is originated from a set of factors that, despite the apparently derangement, act in order to eliminate infection even without the passage of the microorganism to the cerebral tissue.

In this context, NO acts as a key molecule in brain infections. However, it is still unknown if the major problem arises from insufficient concentrations of NO acting directly in the elimination of the parasite, and for this reason, by selecting more resistant strains of the parasite (Gramaglia *et al*. 2006), or if the high concentrations of NO, produced as a result of infection by the protozoan parasite, are responsible for the cerebral edema (Favre *et al*. 1999; Maneerat *et al*. 2000).

In the evaluation of brain oxidative parameters, it was noted an increase in lipid peroxidation for mice treated with dexamethasone in relation to the other groups, mainly in first day post-infection. Nevertheless, the opposite happens with the group of mice treated with L-arginine, where TBARS levels are significantly lower than the other groups (Fig. 4B).

The elevation of TBARS levels for the group treated with dexamethasone may result from a technical artifact, as brain tissue is rich in cholesterol and the drug may form cholesterol hydroperoxides, which may react with thiobarbituric acid, greatly increasing the absorbance of brain samples (Lima and Abdalla 2001). A finding that may corroborate this statement are the dosages of TEAC that do not change in the first days of study for all groups (Fig. 3B). The possibility that lipid peroxidation occurs in this initial period by an increase in IRS was eliminated since uric acid values for this group of animals are similar to those of the other groups until the tenth day of infection (Fig. 6B).

Notwithstanding, it seems that the oxidative effect of nitric oxide was overcome by its vasodilator effect, since the production of uric acid in mice treated with L-arginine was significantly lower when compared to other groups (Fig. 6B), notably from the 10^th^ day of infection. However, the probable vasodilation presented by group L-ARGININE caused no changes on the survival rate of these animals (Fig. 2).

The re-establishment of antioxidant capacity can be decisive for the survival of mice infected with *P. berghei*. The antioxidant capacity decreases significantly in all tested groups. However, only for group DEXAMETHASONE this antioxidant capacity is significantly reversed from the 10^th^ day (Fig. 3B), reinforcing the idea of Favre *et al*. (1999) and Maneerat *et al*. (2000) that the oxidative stress induced by nitric oxide in cerebral microenvironment contributes to the severity of the disease. Another point that deserves to be highlighted for the group treated with dexamethasone is that, despite the inhibition of iNOS, there was only significant increase in serum uric acid concentration from the 15^th^ day (Fig. 6B), signaling that the beginning of IRS coincides with the starting point of deaths in this group. The finding of positive correlation between TBARS and NN for group DEXAMETHASONE corroborates this observation (Additional file Fig. 6).

A factor that may have contributed significantly to the late start of the IRS in this group is the inhibition of the inflammatory process, which is necessary for the occurrence of cytoadherence (Ferreira *et al*. 2004; Pettersson *et al*. 2005). Additionally, this is the only group that displays significant positive correlation between URIC ACID and PARASITEMIA (Additional file Fig. 20), reinforcing the idea that IRS occurs on a temporal scale.

The absence of correlation between NN and PARASITEMIA for all groups and both samples strongly suggests that NO levels do not influence the evolution of parasitemia (Additional file Fig. 9-10).

Considering the different treatments administered, the more promising results were seen with the dexamethasone treatment, since animals exhibited significantly higher survival rate and decreased progression of parasitemia when compared to the other groups. These data suggest that selective inhibition of iNOS, associated to the anti-inflammatory potential of dexamethasone, might decreased lipid peroxidation even with the increase of parasitemia.

In contrast, administration of L-arginine, regardless not significant modification in NN concentrations, promoted vasodilation in both organs, proven by an increase in the concentrations of uric acid, with no effect over the survival rate of these animals.

Nevertheless, the cerebral oxidative changes promoted by the administration of dexamethasone were somehow different from the ones presented by other groups. The re-establishment of the cerebral antioxidant capacity after the 10^th^ day of infection is noteworthy, suggesting the participation of oxidative stress in brain as a result of plasmodial infection, as well as the inhibition of brain NO synthesis, which promoted survival rate of almost 90% of the animals until the 15^th^ day of infection, with possible direct interference of ischemia and reperfusion syndrome, as seen by increased levels of uric acid.

## CONCLUSION

Lately, the role of NO in the physiopathogenesis of malaria has been extensively studied. Nevertheless, its precise involvement in the underlying mechanisms of the disease is still controversial. The present study presents the inhibitory effects of dexamethasone on brain nitric oxide synthesis and its relationship to increased survival in the mice model of malaria. To our best knowledge, it is the first time such results are reported in the scientific literature.

Data of the present study showed that iNOS inhibition by dexamethasone promoted an increase in the survival rate of *P. berghei* -infected animals until the point at which it compromised the functioning of the cerebral microcirculation. Indeed, iNOS inhibition by dexamethasone seems to have stimulated a series of redox effects that, if compensatory hyper stimulated, may be responsible for the worsening of the pulmonary symptoms.

## AUTHOS’ CONTRIBUTION

SP, MDG and MFD were responsible for the design of the study, data analysis, and for the critical revision of the text. DRM, ARQG, ACMGU, MESF, RSS and JRSV were responsible for the collection of data, statistical study and drafting of the manuscript.

## REFERENCES

Adhikari NK, Burns KE, Friedrich JO, Granton JT, Cook DJ and Meade MO (2007) Effect of nitric oxide on oxygenation and mortality in acute lung injury: systematic review and meta-analysis. British Medical Association 334(7597), 779. doi: 10.1136/bmj.39139.716794.55.

Barnes PJ, Adcock I, Spedding M and Vanhoutte PM (1993) Anti-inflammatory actions of steroids: molecular mechanisms. Trends in pharmacological sciences 14(12), 436–441. doi: 10.1016/0165-6147(93)90184-L.

Baron RM, Carvajal IM, Fredenburgh LE, Liu X, Porrata Y, Cullivan ML and Perrella MA (2004) Nitric oxide synthase-2 down-regulates surfactant protein-B expression and enhances endotoxin-induced lung injury in mice. The FASEB Journal 18(11), 1276–1278. doi: 10.1096/fj.04-1518fje.

Becker K, Tilley L, Vennerstrom JL, Roberts D, Rogerson S and Ginsburg H (2004) Oxidative stress in malaria parasite-infected erythrocytes: host–parasite interactions. International Journal for Parasitology 34(2), 163–189. doi: 10.1016/j.ijpara.2003.09.011.

Bein K, Wesselkamper SC, Liu X, Dietsch M, Majumder N, Concel VJ and Borchers MT (2009) Surfactant-associated protein B is critical to survival in nickel-induced injury in mice. American Journal of Respiratory Cell and Molecular Biology 41(2), 226–236. doi: 10.1165/rcmb.2008-0317OC.

Beleslin-Čokić BB, Čokić VP, Wang L, Piknova B, Teng R, Schechter NA and Noguchi CT (2011) Erythropoietin and hypoxia increase erythropoietin receptor and nitric oxide levels in lung microvascular endothelial cells. Cytokine 54(2), 129– 135. doi: 10.1016/j.cyto.2011.01.015.

Botelho C, Silva COS, Beppu OS, Bogossian M and Vargaftig BB (1996) Edema pulmonar na malária: modulaçäo pela cetirizina / Pulmonary edema in murine malaria: modulation by cetirizine. Jornal Brasileiro de Pneumologia 22(2):69–76.

Brinkmann V, Kaufmann SH, Simon MM and Fischer H (1984) Role of macrophages in malaria: O_2_ metabolite production and phagocytosis by splenic macrophages during lethal Plasmodium berghei and self-limiting Plasmodium yoelii infection in mice. Infection and Immunity 44(3), 743–746.

Cabrales P, Zanini GM, Meays D, Frangos JÁ and Carvalho LJ (2011) Nitric oxide protection against murine cerebral malaria is associated with improved cerebral microcirculatory physiology. Journal of Infectious Diseases 203(10), 1454–1463. doi: 10.1093/infdis/jir058.

Chatterjee S, Premachandran S, Shukla J and Poduval TB (2007) Synergistic therapeutic potential of dexamethasone and L-arginine in lipopolysaccharideinduced septic shock. Journal of Surgical Research 140(1), 99–108. doi: 10.1016/j.jss.2006.09.002.

Chiaramonte MG, Donaldson DD, Cheever AW and Wynn TA (1999) An IL-13 inhibitor blocks the development of hepatic fibrosis during a T-helper type 2–dominated inflammatory response. The Journal of Clinical Investigation 104(6), 777–785. doi: 10.1172/JCI7325.

Chiwakata CB, Hemmer CJ and Dietrich M (2000) High levels of inducible nitric oxide synthase mRNA are associated with increased monocyte counts in blood and have a beneficial role in Plasmodium falciparum malaria. Infection and Immunity 68(1), 394–399. doi: 10.1128/IAI.68.1.394-399.2000.

Christofidou-Solomidou M, Scherpereel A, Wiewrodt R, Ng K, Sweitzer T, Arguiri E and Muzykantov VR (2003) PECAM-directed delivery of catalase to endothelium protects against pulmonary vascular oxidative stress. American Journal of Physiology - Lung Cellular and Molecular Physiology 285(2), L283–L292. doi: 10.1152/ajplung.00021.2003.

Cox F (2002) History of human parasitology. Clinical Microbiology Reviews 15:595–612. doi: 10.1128/CMR.15.4.595-612.2002.

De Vera ME, Taylor BS, Wang QI, Shapiro RA, Billiar TR and Geller DA (1997) Dexamethasone suppresses iNOS gene expression by upregulating I-κBα and inhibiting NF-κB. American Journal of Physiology - Gastrointestinal and Liver Physiology 273(6), G1290–G1296. doi: 10.1152/ajpgi.1997.273.6.G1290.

Dell RB, Holleran S, Ramakrishnan R (2002) Sample size determination. Institute of Laboratory Animal Resources Journal 43(4):207–13. doi: 10.1093/ilar.43.4.207.

Dey S, Guha M, Alam A, Goyal M, Bindu S, Pal C and Bandyopadhyay U (2009) Malarial infection develops mitochondrial pathology and mitochondrial oxidative stress to promote hepatocyte apoptosis. Free Radical Biology and Medicine 46(2), 271–281. doi: 10.1016/j.freeradbiomed.2008.10.032.

Dhangadamajhi G, Mohapatra BN, Kar SK and Ranjit M (2009) Endothelial nitric oxide synthase gene polymorphisms and Plasmodium falciparum infection in Indian adults. Infection and Immunity 77(7), 2943–2947. doi: 10.1128/IAI.00083-09.

Di Rosa M, Radomski M, Carnuccio R and Moncada S (1990) Glucocorticoids inhibit the induction of nitric oxide synthase in macrophages. Biochemical and Biophysical Research Communications 172(3), 1246–1252.

Dondorp AM, Omodeo-Salé F, Chotivanich K, Taramelli D and White NJ (2003) Oxidative stress and rheology in severe malaria. Redox report 8(5), 292–294. doi: 10.1179/135100003225002934.

Farombi EO, Shyntum YY and Emerole GO (2003) Influence of chloroquine treatment and Plasmodium falciparum malaria infection on some enzymatic and non-enzymatic antioxidant defense indices in humans. Drug and Chemical Toxicology 26(1), 59–71. doi: 10.1081/DCT-120017558.

Favre N, Ryffel B and Rudin W (1999) The development of murine cerebral malaria does not require nitric oxide production. Parasitology 118(2), 135–138. doi: 10.1016/S1286-4579(99)80513-9.

Ferreira MU, da Silva Nunes M and Wunderlich G (2004) Antigenic diversity and immune evasion by malaria parasites. Clinical and Diagnostic Laboratory Immunology 11(6), 987–995. doi: 10.1128/CDLI.11.6.987-995.2004.

Förstermann U and Sessa WC (2012) Nitric oxide synthases: regulation and function. European Heart Journal 33(7), 829–837. doi: 10.1093/eurheartj/ehr304.

Gachot B, Wolff M, Nissack G, Veber B and Vachon F (1995) Acute lung injury complicating imported Plasmodium falciparum malaria. Chest 108(3), 746–749. doi: 10.1378/chest.108.3.746.

Gaston B, Drazen JM, Loscalzo J and Stamler JS (1994) The biology of nitrogen oxides in the airways. American Journal of Respiratory and Critical Care Medicine 149(2), 538–551. doi: 10.1164/ajrccm.149.2.7508323.

Geller DA, Freeswick PD, Nguyen D, Nussler AK, Di Silvio M, Shapiro RA and Billiar TR (1994) Differential induction of nitric oxide synthase in hepatocytes during endotoxemia and the acute-phase response. Archives of Surgery 129(2), 165–171. doi: 10.1001/archsurg.1994.01420260061008.

Gilbert RS and Herschman HR (1993) “Macrophage” nitric oxide synthase is a glucocorticoid-inhibitable primary response gene in 3T3 cells. Journal of Cellular Physiology 157(1), 128–132. doi: 10.1002/jcp.1041570117.

Gillrie MR, Krishnegowda G, Lee K, Buret AG, Robbins SM, Looareesuwan S and Ho M (2007) Src-family kinase–dependent disruption of endothelial barrier function by Plasmodium falciparum merozoite proteins. Blood 110(9), 3426–3435. doi: 10.1182/blood-2007-04-084582.

Gomes BAQ, da Silva LF, Gomes ARQ, Moreira DR, Dolabela MF, Santos RS and Percário S (2015) N-acetyl cysteine and mushroom Agaricus sylvaticus supplementation decreased parasitaemia and pulmonary oxidative stress in a mice model of malaria. Malaria Journal 14(1), 202. doi: 10.1186/s12936-015-0717-0.

Gramaglia I, Sobolewski P, Meays D, Contreras R, Nolan JP, Frangos JA and Van Der Heyde HC (2006) Low nitric oxide bioavailability contributes to the genesis of experimental cerebral malaria. Nature Medicine 12(12), 1417. doi: 10.1038/nm1499.

Halliwell B and Gutteridge JM (2015) Free radicals in biology and medicine. 5ed. Oxford University Press, USA.

Herrera-Ortíz A, Lanz-Mendoza H, Martínez-Barnetche J, Hernández-Martínez S, Villarreal-Treviño C, Aguilar-Marcelino L and Rodríguez MH (2004) Plasmodium berghei ookinetes induce nitric oxide production in Anopheles pseudopunctipennis midguts cultured in vitro. Insect Biochemistry and Molecular Biology 34(9), 893– 901. doi: 10.1016/j.ibmb.2004.05.007.

Hesse M, Cheever AW, Jankovic D and Wynn TA (2000) NOS-2 mediates the protective anti-inflammatory and antifibrotic effects of the Th1-inducing adjuvant, IL-12, in a Th2 model of granulomatous disease. The American Journal of Pathology 157(3), 945–955. doi: 10.1016/S0002-9440(10)64607-X.

Huber SM, Uhlemann AC, Gamper NL, Duranton C, Kremsner PG and Lang F (2002) Plasmodium falciparum activates endogenous Cl− channels of human erythrocytes by membrane oxidation. The EMBO Journal 21(1-2), 22–30. doi: 10.1093/emboj/21.1.22.

Jaramillo M, Godbout M and Olivier M (2005) Hemozoin induces macrophage chemokine expression through oxidative stress-dependent and-independent mechanisms. The Journal of Immunology 174(1), 475–484. doi: 10.4049/jimmunol.174.1.475.

Jaramillo M, Gowda DC, Radzioch D and Olivier M (2003) Hemozoin increases IFN-γ-inducible macrophage nitric oxide generation through extracellular signal-regulated kinase-and NF-κB dependent pathways. The Journal of Immunology 171(8), 4243–4253. doi: 10.4049/jimmunol.171.8.4243.

Katzung BG and Trevor AJ (2017). Farmacologia Básica e Clínica. 13ed. McGraw Hill, Brasil.

Keller CC, Kremsner PG, Hittner JB, Misukonis MA, Weinberg JB and Perkins DJ (2004) Elevated nitric oxide production in children with malarial anemia: hemozoin-induced nitric oxide synthase type 2 transcripts and nitric oxide in blood mononuclear cells. Infection and Immunity 72(8), 4868–4873. doi: 10.1128/IAI.72.8.4868-4873.2004.

Kinsella JP, Parker TA, Davis JM and Abman SH (2005) Superoxide dismutase improves gas exchange and pulmonary hemodynamics in premature lambs. American Journal of Respiratory and Critical Care Medicine 172(6), 745–749. doi: 10.1164/rccm.200501-146OC.

Kleinert H, Pautz A, Linker K and Schwarz PM (2004) Regulation of the expression of inducible nitric oxide synthase. European Journal of Pharmacology 500(1-3), 255–266. doi: 10.1016/j.ejphar.2004.07.030.

Kohn HI and Liversedge M (1944) On a new aerobic metabolite whose production by brain is inhibited by apomorphine, emetine, ergotamine, epinephrine, and menadione. Journal of Pharmacology and Experimental Therapeutics 82(3), 292– 300.

Korhonen R, Lahti A, Hämäläinen M, Kankaanranta H and Moilanen E (2002) Dexamethasone inhibits inducible nitric-oxide synthase expression and nitric oxide production by destabilizing mRNA in lipopolysaccharide-treated macrophages. Molecular Pharmacology 62(3), 698–704. doi: 10.1124/mol.62.3.698.

Kumar S and Bandyopadhyay U (2005) Free heme toxicity and its detoxification systems in human. Toxicology Letters 157(3), 175–188. doi: 10.1016/j.toxlet.2005.03.004.

Lacerda MVGD, Mourão MPG, Santos PJTD and Alecrim MDGC (2009) Algid malaria: a syndromic diagnosis. Revista da Sociedade Brasileira de Medicina Tropical 42(1), 79–81.

Lee CG, Homer RJ, Zhu Z, Lanone S, Wang X, Koteliansky V and Senior RM (2001) Interleukin-13 induces tissue fibrosis by selectively stimulating and activating transforming growth factor β1. Journal of Experimental Medicine 194(6), 809–822. doi: 10.1084/jem.194.6.809.

Lima ES and Abdalla DSP (2001) Peroxidação lipídica: mecanismos e avaliação em amostras biológicas/ Lipid Peroxidation: Mechanisms and evaluation in biological samples. Brazilian Journal of Pharmaceutical Sciences 37(3), 293–303.

Luse SA and Miller LH (1971) Plasmodium falciparum malaria. The American Journal of Tropical Medicine and Hygiene 20(5), 655–660. doi: 10.4269/ajtmh.1971.20.655.

Maneerat Y, Viriyavejakul P, Punpoowong B, Jones M, Wilairatana P, Pongponratn E and Udomsangpetch R (2000) Inducible nitric oxide synthase expression is increased in the brain in fatal cerebral malaria. Histopathology 37(3), 269–277. doi: 10.1046/j.1365-2559.2000.00989.x.

McClintock DE, Ware LB, Eisner MD, Wickersham N, Thompson BT and Matthay MA (2007) Higher urine nitric oxide is associated with improved outcomes in patients with acute lung injury. American Journal of Respiratory and Critical Care Medicine 175(3), 256–262. doi:10.1164/rccm.200607-947OC.

Mikawa K, Nishina K, Takao Y and Obara H (2003) ONO-1714, a nitric oxide synthase inhibitor, attenuates endotoxin-induced acute lung injury in rabbits. Anesthesia & Analgesia 97(6), 1751–1755. doi:10.1213/01.ANE.0000086896.90343.13.

Miller NJ, Rice-Evans C, Davies MJ, Gopinathan V and Milner A (1993) A novel method for measuring antioxidant capacity and its application to monitoring the antioxidant status in premature neonates. Clinical Science 84(4), 407–412. doi: 10.1042/cs0840407.

Modolell M, Corraliza IM, Link F, Soler G and Eichmann K (1995) Reciprocal regulation of the nitric oxide synthase/arginase balance in mouse bone marrow- derived macrophages by TH 1 and TH 2 cytokines. European Journal of Immunology 25(4), 1101–1104. doi: 10.1002/eji.1830250436.

Muzaffar S, Jeremy JY, Angelini GD, Stuart-Smith K and Shukla N (2003) Role of the endothelium and nitric oxide synthases in modulating superoxide formation induced by endotoxin and cytokines in porcine pulmonary arteries. Thorax 58(7), 598–604. doi:10.1136/thorax.58.7.598.

Nahrevanian H and Dascombe MJ (2006) Simultaneous increases in immunecompetent cells and nitric oxide in the spleen during Plasmodium berghei infection in mice. Journal of Microbiology Immunology and Infection, 39(1), 11.

Narsaria N, Mohanty C, Das BK, Mishra SP and Prasad R (2012) Oxidative stress in children with severe malaria. Journal of Tropical Pediatrics 58(2), 147–150. doi: 10.1093/tropej/fmr043.

Neill AL and Hunt NH (1995) Effects o endotoxin and dexamethasone on cerebral malaria in mice. Parasitology 111(pt4), 443–454.

Omodeo-Salé F, Motti A, Basilico N, Parapini S, Olliaro P and Taramelli D (2003) Accelerated senescence of human erythrocytes cultured with Plasmodium falciparum. Blood 102(2), 705–711. doi: 10.1182/blood-2002-08-243.

Otto TD, Böhme U, Jackson AP, Hunt M, Franke-Fayard B, Hoeijmakers WA and Cunningham D (2014) A comprehensive evaluation of rodent malaria parasite genomes and gene expression. BMC Biology 12(1), 86. doi: 10.1186/s12915-014- 0086-0.

Oury TD, Schaefer LM, Fattman CL, Choi A, Weck KE and Watkins SC (2002) Depletion of pulmonary EC-SOD after exposure to hyperoxia. American Journal of Physiology-Lung Cellular and Molecular Physiology 283(4), L777–L784. doi: 10.1152/ajplung.00011.2002.

Pabón A, Carmona J, Burgos LC and Blair S (2003) Oxidative stress in patients with non-complicated malaria. Clinical Biochemistry 36(1), 71–78. doi: 10.1016/S0009-9120(02)00423-X.

Penet MF, Kober F, Confort-Gouny S, Le Fur Y, Dalmasso C, Coltel N and Viola A (2007) Magnetic resonance spectroscopy reveals an impaired brain metabolic profile in mice resistant to cerebral malaria infected with Plasmodium berghei ANKA. Journal of Biological Chemistry 282(19), 14505–14514. doi: 10.1074/jbc.M608035200.

Percário S, Moreira DR, Gomes BA, Ferreira ME, Gonçalves ACM, Laurindo PS and Green MD (2012) Oxidative stress in malaria. International Journal of Molecular Sciences 13(12), 16346–16372. doi: 10.3390/ijms131216346.

Percario S, Vital ACC, Jablonka F (1994) Dosagem do malondialdeido/Malondialdehyde measurement. Newslab 2(6):46–50.

Peterson TM, Gow AJ and Luckhart S (2007) Nitric oxide metabolites induced in Anopheles stephensi control malaria parasite infection. Free Radical Biology and Medicine 42(1), 132–142. doi: 10.1016/j.freeradbiomed.2006.10.037.

Pettersson F, Vogt AM, Jonsson C, Mok BW, Shamaei-Tousi A, Bergström S and Wahlgren M (2005) Whole-body imaging of sequestration of Plasmodium falciparum in the rat. Infection and Immunity 73(11), 7736–7746. doi: 10.1128/IAI.73.11.7736-7746.200.

Pfeilschifter J and Schwarzenbach H (1990) Interleukin 1 and tumor necrosis factor stimulate cGMP formation in rat renal mesangial cells. Federation of European Biochemical Societies Letters 273(1-2), 185–187. doi: 10.1016/0014- 5793(90)81080-8.

Planche T, Macallan DC, Sobande T, Borrmann S, Kun JF, Krishna S and Kremsner PG (2010) Nitric oxide generation in children with malaria and the NOS2G-954C promoter polymorphism. American Journal of Physiology-Regulatory, Integrative and Comparative Physiology 299(5), R1248–R1253. doi: 10.1152/ajpregu.00390.2010.

Potter SM, Mitchell AJ, Cowden WB, Sanni LA, Dinauer M, De Haan JB and Hunt NH (2005) Phagocyte-derived reactive oxygen species do not influence the progression of murine blood-stage malaria infections. Infection and Immunity 73(8), 4941–4947. doi: 10.1128/IAI.73.8.4941-4947.2005.

Rabkin DG, Sladen RN, DeMango A, Steinglass KM and Goldstein DJ (2001) Nitric oxide for the treatment of postpneumonectomy pulmonary edema. The Annals of thoracic surgery 72(1), 272–274. doi: 10.1016/S0003-4975(01)02476-6.

Radomski MW, Palmer RM and Moncada S (1990) Glucocorticoids inhibit the expression of an inducible, but not the constitutive, nitric oxide synthase in vascular endothelial cells. Proceedings of the National Academy of Sciences 87(24), 10043– 10047.

Rahman I and MacNee W (2000) Oxidative stress and regulation of glutathione in lung inflammation. European Respiratory Journal 16(3), 534–554. doi: 10.1034/j.1399-3003.2000.016003534.x.

Re R, Pellegrini N, Proteggente A, Pannala A, Yang M and Rice-Evans C (1999) Antioxidant activity applying an improved ABTS radical cation decolorization assay. Free Radical Biology and Medicine 26(9-10), 1231–1237. doi: 10.1016/S0891-5849(98)00315-3.

Saeftel M, Krueger A, Arriens S, Heussler V, Racz P, Fleischer B and Hoerauf A (2004) Mice deficient in interleukin-4 (IL-4) or IL-4 receptor α have higher resistance to sporozoite infection with Plasmodium berghei (ANKA) than do naive wild-type mice. Infection and Immunity 72(1), 322–331. doi: 10.1128/IAI.72.1.322-331.2004.

Saxena S, Kumar R, Madan T, Gupta V, Muralidhar K and Sarma PU (2005) Association of polymorphisms in pulmonary surfactant protein A1 and A2 genes with high-altitude pulmonary edema. Chest 128(3), 1611–1619. doi: 10.1378/chest.128.3.1611.

Sanni LA, Thomas SR, Tattam BN, Moore DE, Chaudhri G, Stocker R and Hunt NH (1998) Dramatic changes in oxidative tryptophan metabolism along the kynurenine pathway in experimental cerebral and noncerebral malaria. American Journal of Pathology 152(2), 611–619.

Schreiber MD, Gin-Mestan K, Marks JD, Huo D, Lee G and Srisuparp P (2003) Inhaled nitric oxide in premature infants with the respiratory distress syndrome. New England Journal of Medicine 349(22), 2099–2107. doi: 10.1056/NEJMoa031154.

Silverthorn DU. Fisiologia humana: uma abordagem integrada/Human physiology: an integrated approach. 5ed. Porto Alegre: Artmed; 2010.

Skimming JW, Nasiroglu O, Huang CJ, Wood CE, Stevens BR, Haque IU and Sarcia PJ (2003) Dexamethasone suppresses iNOS yet induces GTPCH and CAT-2 mRNA expression in rat lungs. American Journal of Physiology-Lung Cellular and Molecular Physiology 285(2), L484–L491. doi: 10.1152/ajplung.00433.2002.

Sohail M, Kaul A, Raziuddin M and Adak T (2007) Decreased glutathione-S-transferase activity: Diagnostic and protective role in vivax malaria. Clinical Biochemistry 40(5-6), 377–382. doi: 10.1016/j.clinbiochem.2007.01.005.

Suresh KP (2011) An overview of randomization techniques: an unbiased assessment of outcome in clinical research. Journal of Human Reproductive Sciences 4(1), 8. doi: 10.4103/0974-1208.82352.

Syarifah HP, Hayano M and Kojima S (2003) Cytokine and chemokine responses in a cerebral malaria-susceptible or-resistant strain of mice to Plasmodium berghei ANKA infection: early chemokine expression in the brain. International Immunology 15(5), 633–640. doi: 10.1093/intimm/dxg065.

Taylor WR, Cañon V and White NJ (2006) Pulmonary manifestations of malaria. Treatments in Respiratory Medicine 5(6), 419–428.

er Horst SA, Walther FJ, Poorthuis BJ, Hiemstra PS and Wagenaar GT (2007) Inhaled nitric oxide attenuates pulmonary inflammation and fibrin deposition and prolongs survival in neonatal hyperoxic lung injury. American Journal of Physiology-Lung Cellular and Molecular Physiology 293(1), L35–L44. doi: 10.1152/ajplung.00381.2006.

Vale VV, Vilhena TC, Trindade RCS, Ferreira MRC, Percário S, Soares LF, Pereira WLA, Brandão GC, Oliveira AB, Dolabela MF and De Vasconcelos F (2015) Anti-malarial activity and toxicity assessment of Himatanthus articulatus, a plant used to treat malaria in the Brazilian Amazon. Malaria Journal 14(1), 132. doi: 10.1186/s12936-015-0643-1.

Van Der Heyde HC, Gu Y, Zhang Q, Sun G and Grisham MB (2000) Nitric oxide is neither necessary nor sufficient for resolution of Plasmodium chabaudi malaria in mice. The Journal of Immunology 165:3317–23. doi: 10.4049/jimmunol.165.6.3317.

Walker G, Pfeilschifter J and Kunz D (1997) Mechanisms of suppression of inducible nitric-oxide synthase (iNOS) expression in interferon (IFN)-γ-stimulated RAW 264.7 cells by dexamethasone: evidence for glucocorticoid-induced degradation of iNOS protein by calpain as a key step in post-transcriptional regulation. Journal of Biological Chemistry 272(26), 16679–16687. doi: 10.1074/jbc.272.26.16679.

Wedgwood S, Lakshminrusimha S, Farrow KN, Czech L, Gugino SF, Soares F and Steinhorn RH (2012) Apocynin improves oxygenation and increases eNOS in persistent pulmonary hypertension of the newborn. American Journal of Physiology-Lung Cellular and Molecular Physiology 302(6), L616–L626. doi: 10.1152/ajplung.00064.2011.

Wilmanski J, Siddiqi M, Deitch EA and Spolarics Z (2005) Augmented IL-10 production and redox-dependent signaling pathways in glucose-6-phosphate dehydrogenase-deficient mouse peritoneal macrophages. Journal of leukocyte biology 78(1), 85. doi: 10.1189/jlb.0105010.

World Health Organization. WHO Global Malaria Program. World Malaria Report 2017. https://www.who.int/malaria/publications/world-malaria-report-2017/en/. Accessed 08 December 2018.

World Health Organization. WHO Global Malaria Program. World Malaria Report 2014. http://www.who.int/malaria/publications/world_malaria_report_2014/en/. Accessed 16 December 2015.

World Health Organization. WHO Global Malaria Program. World Malaria Report 2011. http://www.who.int/malaria/publications/atoz/978924156. Accessed 21 March 2013.

Yazar S, Kilic E, Saraymen R and Ozbilge H (2004) Serum malondialdehyde levels in patients infected with Plasmodium vivax. The West Indian Medical Journal 53(3), 147–149.

Yeo TW, Lampah DA, Gitawati R, Tjitra E, Kenangalem E, McNeil YR and Price RN (2007) Impaired nitric oxide bioavailability and L-arginine–reversible endothelial dysfunction in adults with falciparum malaria. Journal of Experimental Medicine 204(11), 2693–2704. doi: 10.1084/jem.20070819.

Yeo TW, Lampah DA, Gitawati R, Tjitra E, Kenangalem E, Piera K and Anstey NM (2008) Angiopoietin-2 is associated with decreased endothelial nitric oxide and poor clinical outcome in severe falciparum malaria. Proceedings of the National Academy of Sciences 105(44), 17097–17102. doi: 10.1073/pnas.0805782105.

